# The effects of the NMDAR co-agonist D-serine on the structure and function of the optic tectum

**DOI:** 10.1101/2021.08.25.457610

**Authors:** Zahraa Chorghay, Vanessa J. Li, Arna Ghosh, Anne Schohl, Edward S. Ruthazer

## Abstract

The N-methyl-D-aspartate type glutamate receptor (NMDAR) is a molecular coincidence detector which converts correlated patterns of neuronal activity into cues for the structural and functional refinement of developing circuits in the brain. D-serine is an endogenous co-agonist of the NMDAR. In this study, we investigated the effects of potent enhancement of NMDAR-mediated currents by chronic administration of saturating levels of D-serine on the developing *Xenopus* retinotectal circuit. Chronic exposure to the NMDAR co-agonist D-serine resulted in structural and functional changes to the optic tectum. D-serine administration affected synaptogenesis and dendritic morphology in recently differentiated tectal neurons, resulting in increased arbor compaction, reduced branch dynamics, and higher synapse density. These effects were not observed in more mature neurons. Calcium imaging to examine retinotopic map organization revealed that tectal neurons of animals raised in D-serine had sharper visual receptive fields. These findings suggest that the availability of endogenous NMDAR co-agonists like D-serine at glutamatergic synapses may regulate the refinement of circuits in the developing brain.

**Significance statement:** N-methyl-D-aspartate receptors (NMDAR) are implicated in activity-dependent circuit plasticity. We used administration of the NMDAR co-agonist D-serine to further examine the role of the NMDAR in circuit development *in vivo*. D-serine stabilised dendritic arbors specifically of recently differentiated neurons, promoted synaptogenesis, and led to sharper retinotopic receptive fields in the optic tectum. Together, these results support the idea that signaling in response to synaptic current through NMDARs promotes the maturation of developing brain circuits.

## Introduction

During the development of functional circuits, neuronal processes elaborate and establish coarse topographic maps, then undergo synaptic and structural refinement to enable precise connectivity. The N-methyl-D-aspartate type glutamate receptor (NMDAR) appears to play an evolutionarily conserved role in the activity-dependent selection of inputs for refinement (Ewald and Cline, 2009). While NMDARs are heterogenous in their composition, they classically require simultaneous ligand binding of glutamate and a co-agonist, either glycine or D-serine (Wolosker, 2007), as well as sufficient depolarization to relieve the magnesium block of the ion channel pore (Mayer et al., 1984; Nowak et al., 1984). The concurrent ligand-binding and depolarizing potential requirements for receptor activation make NMDARs ideal for detection of coincident presynaptic glutamate release and postsynaptic depolarization (Bliss and Collingridge, 1993).

This suggests a model whereby NMDAR activation converts patterned neuronal activity into downstream signaling cascades that direct the refinement of topographic maps. Correlated activity is thought to mediate synaptic strengthening and promote arbor stabilization, prolonging branch lifetimes, and suppressing branch dynamics (Kutsarova et al., 2016). Conversely, uncorrelated activity promotes branch destabilization, including increased branch addition, loss, and elongation. Indeed, loss of NMDAR function perturbs in arbor growth and branch dynamics of both axons and dendrites (Rocha and Sur, 1995; Rajan and Cline, 1998; Rajan et al., 1999; Sin et al., 2002; Ruthazer et al., 2003; Munz et al., 2014), leading to disorganization of afferent projections during the development of topographic maps (Cline et al., 1987; Cline and Constantine-Paton, 1989; Simon et al., 1992; Iwasato et al., 1997; Lee et al., 2005a; Lee et al., 2005b; Zhou et al., 2021).

Furthermore, NMDARs have been found to differentially affect morphology, plasticity and circuit function depending upon their presynaptic or postsynaptic localization (Corlew et al., 2008; Kesner et al., 2020).

D-serine is found endogenously in the brain in a similar distribution to that of NMDARs (Hashimoto et al., 1993; Schell et al., 1997) and enhances NMDAR-dependent synaptic transmission (Mothet et al., 2000; Papouin et al., 2012; Rosenberg et al., 2013). D-serine has been implicated in hippocampal long-term potentiation (Yang et al., 2003; Henneberger et al., 2010; Rosenberg et al., 2013; Han et al., 2015) and depression (Duffy et al., 2008), spatial reversal learning, extinction of fear conditioning (Labrie et al., 2009), and facilitating representation of sequences (DeVito et al., 2011). In the nervous system, whether the primary source of D-serine is glia (Papouin et al., 2017) or neurons (Kartvelishvily et al., 2006; Rosenberg et al., 2010) remains controversial, and likely depends on the brain region, developmental stage, and presence of pathology (Ivanov and Mothet, 2019; Coyle et al., 2020).

The role of NMDARs in developmental plasticity has largely been characterized through loss-of-function manipulations, which lack specificity, as they disrupt normal network activity by reducing overall neuronal excitation. The NMDAR co-agonist glycine has a role beyond its function as a NMDAR co-ligand in the central nervous system as a major inhibitory neurotransmitter, directly affecting circuit excitability. In contrast, administration of D-serine is a more NMDAR-specific pharmacological manipulation that enhances basal NMDAR currents while preserving other aspects of ongoing glutamatergic neurotransmission. Therefore, we used D-serine administration as a gain-of-function manipulation to study the effects of NMDAR-specific signal enhancement on circuit development. In the *Xenopus* tadpole visual system, where retinal ganglion cell (RGC) axons project contralaterally to the optic tectum, D-serine has been reported to promote the maturation of glutamatergic synapses, stabilize axonal arbor structure, and enhance functional inputs of RGCs onto tectal cells through NMDAR-dependent mechanisms (Van Horn et al., 2017). Here, we further investigated the role of enhanced NMDAR activity in circuit refinement by studying the effects of D-serine on the structure and function of postsynaptic neurons in the optic tectum. We found that D-serine administration led to compaction of dendritic arbor morphology in recently differentiated tectal neurons, increased synaptic density, and sharpened receptive fields in the optic tectum.

## Materials and Methods

### Husbandry and Animals

All procedures were approved by the Animal Care Committee of the Montreal Neurological Institute at McGill University in accordance with Canadian Council on Animal Care guidelines. For generation of tadpoles, female adult albino *Xenopus laevis* frogs (RRID:XEP_Xla300) from our in-house breeding colony were primed with 50 IU pregnant mare serum gonadotropin (PMSG; Prospec Bio HOR-272), and 3 days later, were injected with 400 IU human chorionic gonadotropin (hCG; Sigma-Aldrich CG10; RRID:SCR_018232).

### Natural fertilization (Fig 1, 2, 3)

Males were injected with 150 IU hCG on the same day as females, and placed together to induce amplexus. Eggs were collected the following day.

**Figure 1.**
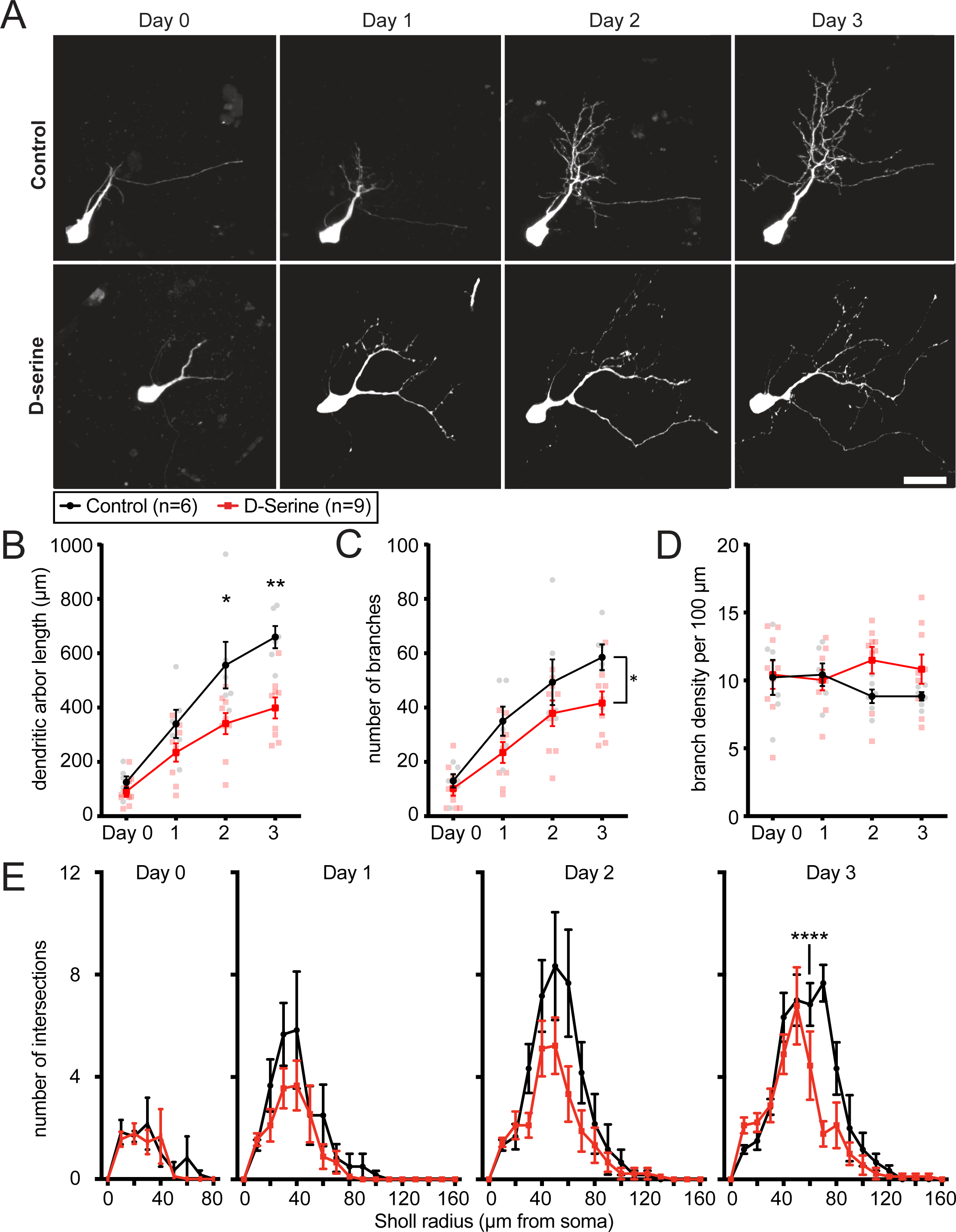
Growth of dendritic arbors in recently differentiated tectal cells of animals reared and imaged in D-serine over 4 d. (A) Representative images. Scale bar: 20 µm. For animals reared in D-serine compared to control, (B) total dendritic arbor length (Day 1–3 RM ANOVA interaction *p=0.0428, Šídák’s multiple comparisons post-hoc test Day 2 **p=0.0089, Day 3 **p=0.0013) and (C) branch number (Day 1–3 RM ANOVA main effects of time p<0.0001 and treatment *p=0.0397) were reduced, (D) resulting in increased branch density (Day 1–3 RM ANOVA interaction *p=0.0106). (E) Sholl analysis shows tectal dendritic arbors of animals reared in D-serine remain closer to the cell body (RM ANOVA Day 3 corrected interaction ***p=0.0003, Šídák’s post-hoc test for multiple comparisons radius 70 µm ****p<0.0001). [n=9 cells for D-serine, n=6 for control.]

**Figure 2.**
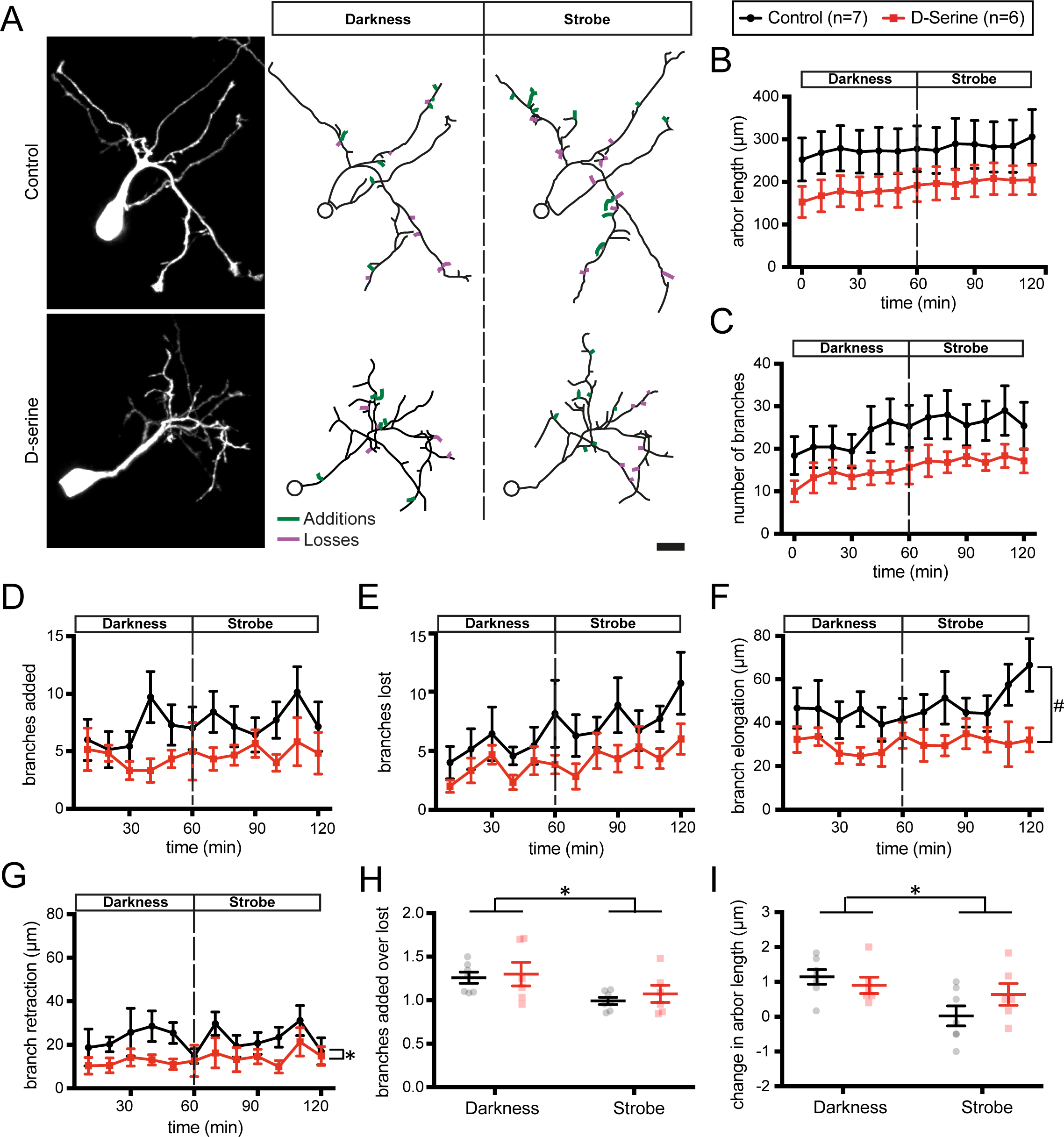
Branch dynamics of tectal cell dendritic arbors in animals reared in D-serine for 48 h. Cells are imaged every 10 mins for 2 h, with 1 h darkness and 1 h of 1 Hz strobe visual stimulation. (A) Representative images. Reconstructed arbors show added (green) and lost (purple) branches between the last 2 time points during darkness and strobe. Scale bar: 10 µm. (B) Total dendritic arbor length (C) branch number, number of dendritic branches (D) added and (E) lost over time were not significantly different between D-serine and control conditions [RM ANOVA p = 0.209, 0.152, 0.167, 0.143 respectively for main effects of treatment]. (F) Branch elongations [RM ANOVA main effect of treatment #p=0.0706] and (G) branch retractions [RM ANOVA main effect of treatment *p=0.038] were reduced following D-serine rearing, compared to control. (H) The ratio of branch additions to subtractions binned by the hour shows a shift from net addition to more equal rates of addition and subtractions under 1 Hz strobe [RM ANOVA main effect of vis stim *p =0.0485]. (I) Changes in arbor length are decreased under 1 Hz strobe [RM ANOVA main effect of vis stim *p=0.0430]. [1 cell per animal analyzed with n=6 cells for D-serine, n=7 for control.]

### In vitro fertilization with GCaMP6 sperm (Fig 3-1)

Eggs from primed females were collected and fertilized *in vitro* using thawed sperm aliquots from transgenic Xla.Tg(tubb2b:GCaMP6s,Rno.elas:GFP)^NXR^ frogs (NXR_0107, Xenopus National Resource, Woods Hole) to generate animals with neuronal GCaMP6s expression.

**Figure 3.**
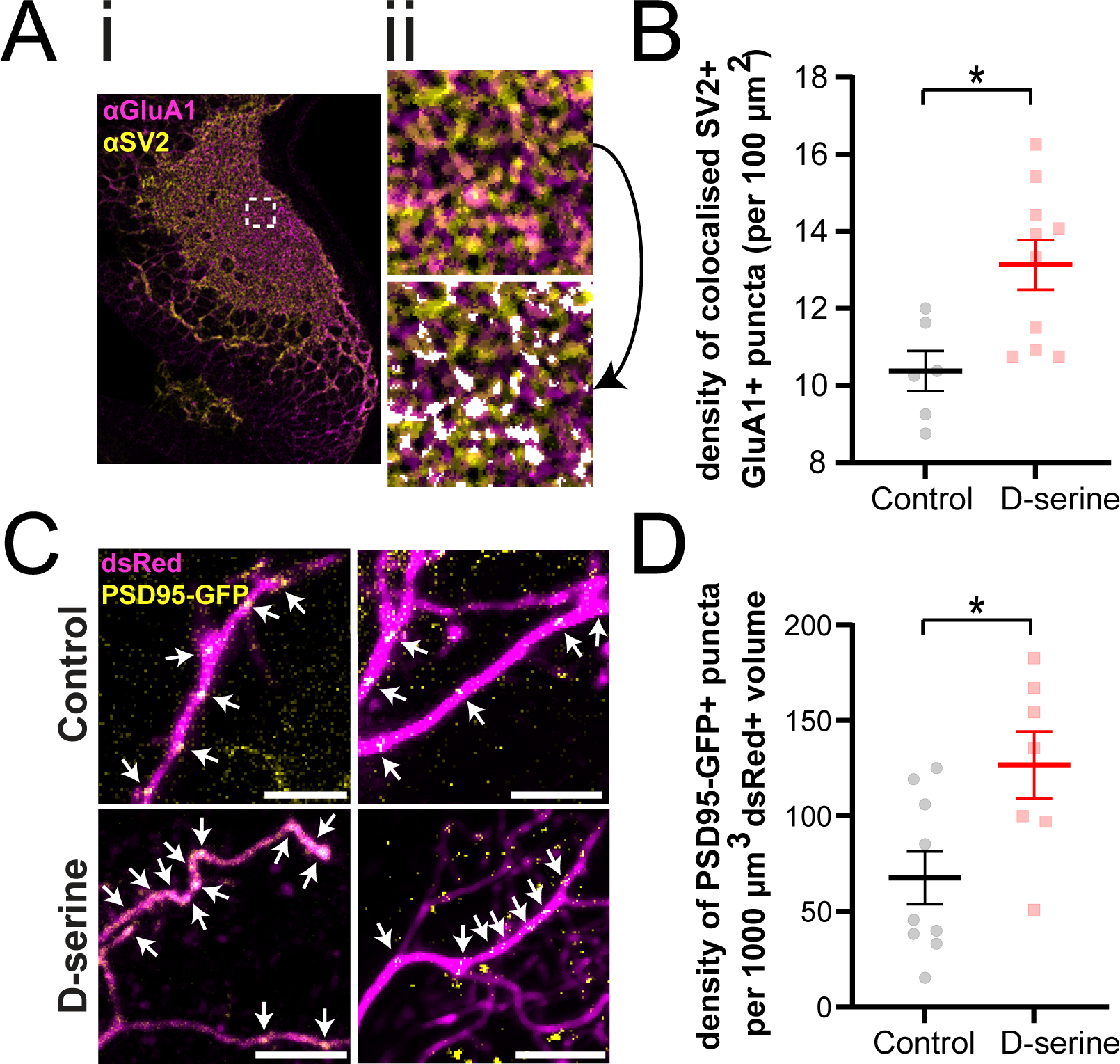
Synaptic density in the tectum of animals reared in D-serine for 48 h. (A) Anatomical synapse density in the tectum was measured by counting colocalized puncta of SV2 (presynaptic) and GluA1 (postsynaptic) immunofluorescence on brain sections of animals reared in D-serine compared to control. For a sample field, (i) the tectum is shown, (ii) zoomed into the dashed inset of an example 20 µm x 20 µm field shown before (top) and after (bottom) automated processing to identify “anatomical synapses” (white overlay). (B) Anatomical synapse density in the optic tectum was significantly different for animals reared in D-serine compared to control medium. [t-test *p=0.0101, 2-3 fields per tectal hemisphere were analysed i.e. at least 5 fields per animal from n=5 animals for D-serine, n=3 for control. Only colocalized puncta that fit the synapse size criterion of 0.1–5.0 µm^2^ were included.] (C) Synapse density along the dendritic arbors of single tectal neurons was quantified by counting PSD95-GFP+ puncta (postsynaptic density) on dsRed+ cells following tectal electroporations and measuring punctum number per dsRed volume. Scale bar: 20 µm. Arrows: examples of puncta. (D) There were significantly more PSD95 puncta on the dendritic arbors of single tectal neurons following rearing in D-serine versus control medium. [t-test *p=0.0176, 1 cell per animal analyzed with n=7 cells for D-serine, n=9 for control.]

### In vitro fertilization with mRNA injection (Fig 4)

Eggs from primed females were collected for *in vitro* fertilization with sperm from male albino frogs and microinjection of GCaMP6s and mCherry messenger ribonucleic acid (mRNA) into one blastomere of two-cell stage embryos as previously described (Kesner et al., 2020). Briefly, a mixture of purified GCaMP6s (500 pg) and mCherry (250 pg) mRNA in 2 nL RNAase-free water was pressure injected into one blastomere of two-cell stage embryos using a calibrated glass micropipette attached to a PLI-100 picoinjector (Harvard Apparatus). Several days after injection, we screened for animals with unilateral mCherry and high levels of GCaMP6s fluorescence for use in calcium imaging experiments. Since RGC axons project contralaterally to the optic tectum, GCaMP6s labels RGC axon terminals and postsynaptic tectal cells in opposite hemispheres of these animals, allowing us to perform calcium imaging of either the RGCs or the tectal cells in each tectal lobe.

**Figure 4.**
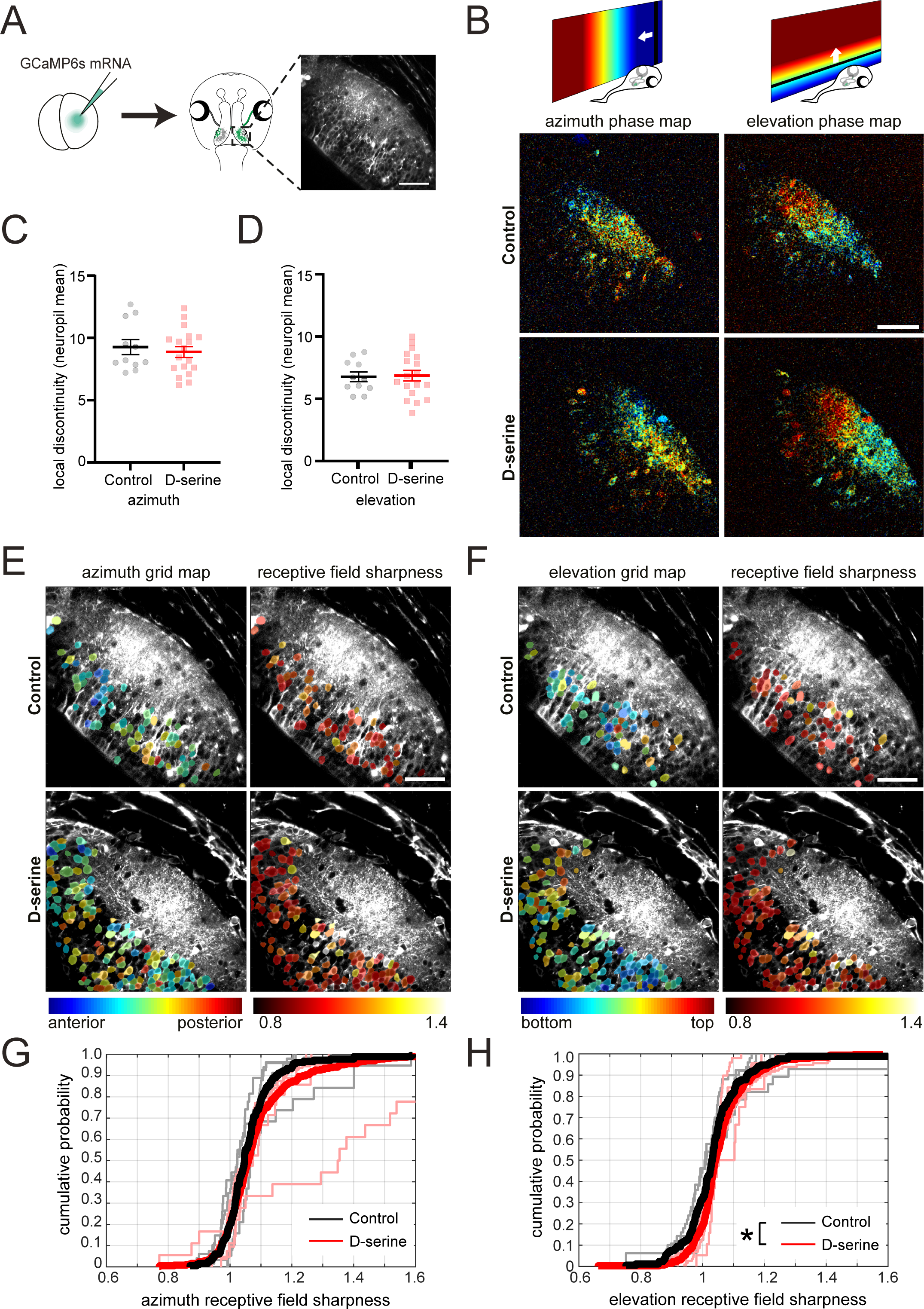
Post-synaptic retinotopic maps in stage 48 animals reared in D-serine from stage 37. (A) mRNA for GCaMP6s was injected into one blastomere at the two-cell stage to generate tadpoles with bilateral mosaic GCaMP6s expression restricted to half the animal. We performed two-photon imaging to observe calcium fluorescence in the optic tectum. Scale bar: 50 µm. (B) Representative images of topographic phase maps in response to drifting bar stimuli in the azimuth and elevation axes from animals reared in D-serine or control medium. The schematics indicate the dynamic range corresponding with the location of the drifting bar in the tadpole’s visual field, serving as the color code for the retinotectal maps. Scale bar: 50 µm. (C–D) Map discontinuity in the (C) azimuth and (D) elevation showed no difference across conditions. [Mann-Whitney test, n=18 animals for D-serine, n=11 for control.] (E-F) Representative images of topographic (left panels) grid maps and (right panels) receptive field sharpness of individual tectal cells in response to a bar flashed at each of 5 locations across the (E) azimuth and (F) elevation axes. The grid maps are color-coded by optimal stimulus position. Scale bar: 50 µm. (G–H) Cumulative distributions of receptive field sharpness of tectal cell bodies in individual animals (lighter lines) and across treatments (darker lines) for the (G) azimuth [K-S Test p=0.1002, n=6 animals and n=365 cells for D-serine, n=5 animals and n=181 cells for control] and (H) elevation axes [K-S Test *p=0.0210, n=6 animals and n=326 cells for D-serine, n=5 animals and n=172 cells for control].

### Tadpole rearing

For all experiments, tadpoles were reared in glass bowls kept in biological oxygen demand incubators set to 21°C with a 12h:12h light:dark cycle. Rearing medium was 0.1X Modified Barth’s Solution with HEPES (MBSH; 88 mM NaCl, 1 mM KCl, 2.4 mM NaHCO3, 0.82 mM MgSO4 x 7H2O, 0.33 mM Ca(NO3)2 x 4H2O, 0.41 mM CaCl2, 10 mM HEPES, pH 7.4).

### Constructs

pCAG-Cre, pCALNL-EGFP, pCALNL-DsRed are a generous gift from CL Cepko and are currently available through Addgene (plasmids 13775, 13770, 13769). All plasmids were grown in DH5a competent cells (Life Technologies) and purified using endotoxin-free maxiprep kits (Qiagen).

### mRNA synthesis

To synthesize the mRNA for blastomere injections, GCaMP6s (Addgene plasmid 40753) and mCherry (plasmid gift of K. Murai) were each cloned into the pCS2+ vector. The GCaMP6s plasmid was cut with NotI / Klenow fill in / BglII, the mCherry plasmid was cut with BamHI / EcoRV, and the pCS2+ vector was cut with BamH1 / SnaB1. For mRNA synthesis, the plasmids in the pCS2+ vector were linearized with NotI, and the capped mRNA of GCaMP6s and mCherry were transcribed with the SP6 mMessage mMachine Kit (Ambion, Thermo Fisher).

### Electroporation (Figs. 1, 2, 3)

Tadpoles at stage 42-44 were anesthetized in MS222 (0.02% in 0.1% MBSH) and placed on a Kimwipe under a dissecting microscope. Cre-mediated single-cell labelling by electroporation (CREMSCLE) for high-efficiency, sparse labelling of the optic tectum was performed (Schohl et al., 2020). A pair of custom-made platinum plate electrodes, connected to an electrical stimulator (SD 9, Grass Instruments), was placed on each side of the brain to deliver current pulses: 38 V, 2-3 ms, 2 pulses at reverse polarity 1 s apart. A 3 µF capacitor was connected in parallel to generate an exponential decay current pulse. For daily imaging (Fig 1), we co-electroporated Cre-recombinase and Cre-dependent EGFP at a ratio of 1:4000 (2.5*10^-4^ µg/µl pCAG-Cre, 1 µg/µl pCALNL-EGFP) injected intraventricularly together with Fast Green dye for visualization. To count synapses per cell (Fig 3C–D), we co-electroporated Cre-recombinase and Cre-dependent dsRed at a ratio of 1:4000 (2.5*10^-4^ µg/µl pCAG-Cre, 1 µg/µl pCALNL-dsRed) and pPSD95-GFP (1 µg/µl).

While CREMSCLE was used to target more immature neurons, we used single-cell electroporation (Haas et al., 2001) to study the morphology of randomly labelled tectal cells. A borosilicate glass micropipette (Sutter Instruments) containing the plasmid (1 µg/µl EGFP) was gently introduced into the brain of anesthetized stage 44–45 tadpoles. Platinum plate electrodes were placed on each side of the brain and a 50 V train of 1 ms pulses at 200 Hz was applied for 0.5 s through the micropipette, with pulse trains repeated twice to ensure delivery of the plasmid.

48 h after electroporation, animals were screened for brightly labelled well-separated tectal cells, then reared in control rearing medium or medium supplemented with 100 µM D-serine (Tocris).

### In vivo two-photon imaging for morphology (Fig 1-3)

Excitation light at 910 nm (EGFP, GCaMP6s) or 990 nm (dsRed) was produced by a Maitai-BB Ti:Sapphire or an InSightX3 femtosecond pulsed IR laser (Spectra Physics).

### Daily imaging (Fig 1, 1-1)

Animals were screened for bright, sparse EGFP expression in the optic tectum and were anesthetized in MS222 (0.02% in 0.1X MBSH), placed in a custom-made Sylgard chamber that fit the tadpole’s body, and placed under a cover glass. Daily z-series images of single tectal neurons were acquired on an Olympus FV300 microscope custom-built for multiphoton imaging with an Olympus LUMPLAFLN 60X water-immersion objective (1.0 NA). Z-series optical sections were collected at 1 µm intervals using Fluoview software (version 5.0). After imaging, the animals were placed in an isolated well of a 6-well plate in D-serine or control rearing medium. Images were collected every day for 4 d, with the animals returned to their respective well and the rearing media changed daily. All image z-stacks were denoised using CANDLE non-local means denoising software implemented in MATLAB (MathWorks) (Coupé et al., 2012), and the three-dimensional reconstruction of single neurons performed using Imaris 6.0 (Bitplane). The neurons analyzed for the control groups (labelled by CREMSCLE in Fig 1, and by single cell electroporation in Fig 1-1) have been previously compared in our CREMSCLE methods publication (Schohl et al., 2020).

**Figure 1-1.**
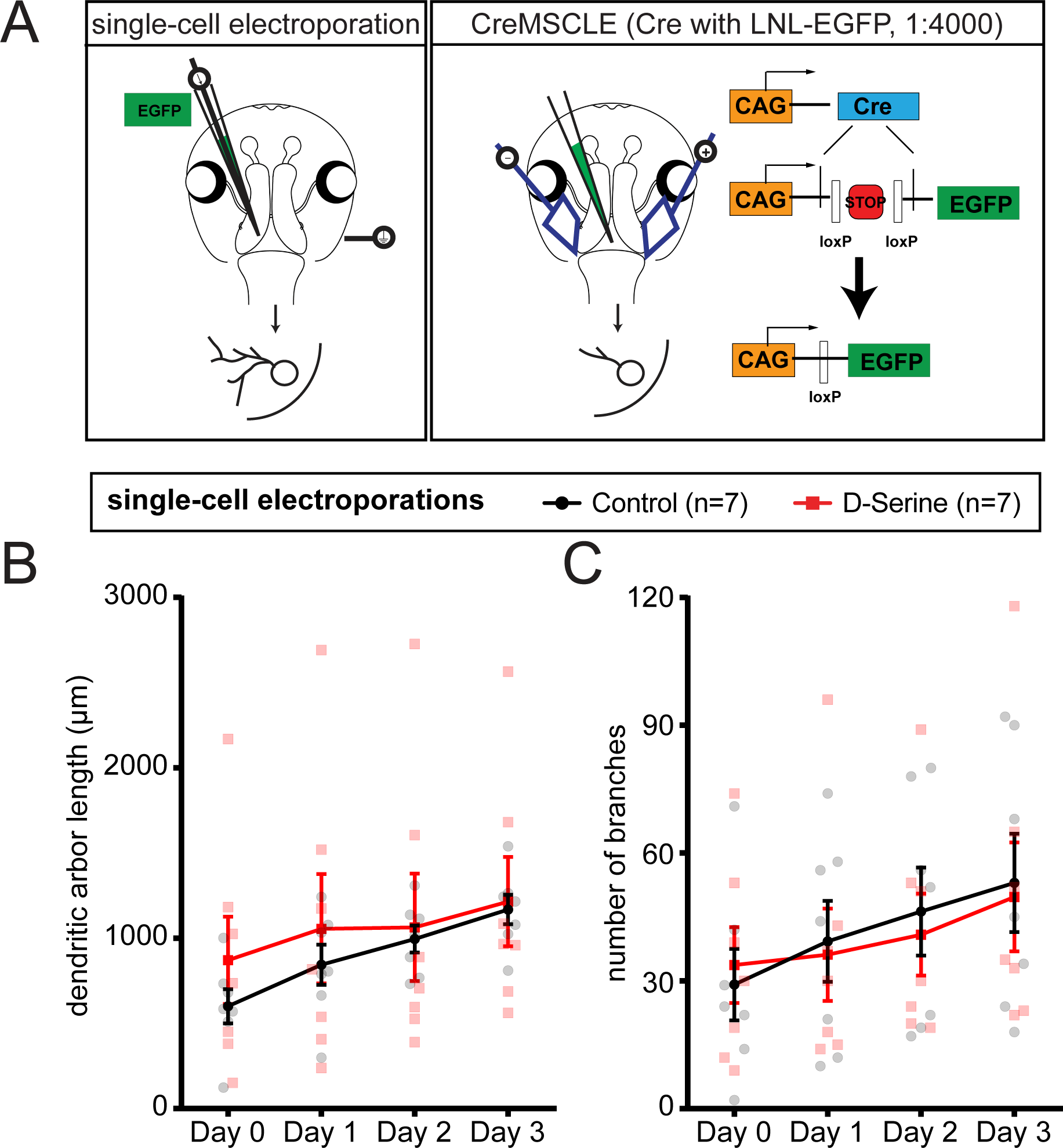
Growth of dendritic arbors in tectal cells labelled by single-cell tectal electroporation of EGFP of animals reared and imaged in D-serine over 4 days. (A) Schematic showing single-cell tectal electroporation versus CREMSCLE. Compared to recently differentiated cells labelled by CREMSCLE, the more mature neurons labelled by single-cell electroporation do not show significant changes in D-serine versus control conditions for the (B) total dendritic arbor length and (C) branch number. [1 cell per animal analyzed with n=7 cells for D-serine, n=7 for control.]

### Short interval imaging (Fig 2)

Animals selected for bright sparse EGFP expression in the optic tectum were placed in isolated wells containing D-serine or control rearing medium. 24 h later, the rearing medium was refreshed, and 48 h after screening (corresponding to day 2 of daily imaging), animals were imaged. Tadpoles were immobilized in 2 mM pancuronium bromide (Tocris) for 3–5 mins, embedded in 0.8% w/v UltraPureTM low melting point agarose on a petri dish, and the petri dish filled with rearing medium. Animals were allowed to stabilize in the imaging set-up for 20 min in darkness prior to beginning each imaging experiment.

Dendritic arbors were imaged with an Olympus XLUMPlanFLN 20X water-immersion objective (1.0 NA) mounted on a commercial high-speed resonance scanner-based multiphoton microscope (Thorlabs) with piezoelectric objective focusing (Physik Instrumente). Z-series optical sections were collected at 1 µm intervals using ThorImage software (version 3.0+). Images were collected every 10 min for 2 h, the first hour in darkness and the second hour with 1 Hz stroboscopic stimulation presented with 10 ms full-field light flashes from a red luxeon LED controlled by a UnoR3 board and a custom Matlab script. All image z-stacks were denoised with CANDLE, and the four-dimensional manual reconstruction for dynamic morphometric analysis performed using Dynamo software implemented in MATLAB (Hossain et al., 2012). Only branches thresholded for a minimum lifetime length of 1.5 µm were included in the quantitative analysis.

### Immunohistochemistry for anatomical synapses (Fig 3A,B)

Animals were anesthetized in 0.02% MS-222, fixed by immersion in 4% paraformaldehyde (Cedarlane/EMS 15735-30-S) in phosphate buffered saline (PBS) for 1 h at room temperature, transferred to ice-cold 100% methanol, and post-fixed overnight at –20°C. Samples were then washed for 1 h in a solution of 100 mM Tris/HCl, pH7.4 with 100 mM NaCl. For infiltration and cryoprotection, the samples were incubated overnight at room temperature in a solution of 15% fish gelatin (Norland HP-03) with 15% sucrose, and subsequently in 25% fish gelatin with 15% sucrose. Samples were embedded and frozen in a solution of 20% fish gelatin with 15% sucrose for cryosectioning. Horizontal sections were collected at 20 µm thickness on a cryostat and directly mounted onto Superfrost-plus slides (Fisher).

Sections were exposed to antigen retrieval with 1% sodium dodecyl sulfate (SDS) in PBS for 3 min at room temperature. Slides were incubated with blocking solution (10% bovine serum albumin and 5% normal goat serum in PBS). Sections were immunostained with rabbit anti-GluA1 (1:200; Abcam ab109450 RRID:AB_10860361) and mouse anti-SV2 (1:1000; Developmental Studies Hybridoma Bank sv2-2a RRID:AB_2315387), and secondary antibodies Alexa-555 goat anti-rabbit IgG (1:200; Invitrogen A-21428 RRID:AB_141784) and Alexa-647 goat anti-mouse IgG (1:200; Invitrogen A21236 RRID:AB_2535805). Confocal images of the optic tectum were acquired with a 20x/0.75 CS2 objective on a Leica SP8 confocal microscope.

#### Synapse quantification (Fig 3A,B)

For synapse quantification, a similar approach to Sild et al. (2016) was used. Briefly, analysis was performed on the optical section 3 μm beneath the cut surface of the histological section in fields devoid of excess vasculature or sectioning irregularities. 20 μm x 20 μm fields were selected from the neuropil region from each imaged tectal hemisphere. Images were pre-processed using FIJI with background subtraction (10 px radius rolling ball) then median filtering (2 px radius), followed by Moments auto-thresholding for each channel. To identify synapses, the logical AND of the SV2 (presynaptic) and GluA1 (postsynaptic) channels was taken, then the Analyze Particles function with the size criterion of 0.1–5.0 μm^2^ area was applied, resulting in an estimate of puncta with pre- and postsynaptic labelling. Analysis was performed blind to experimental condition.

#### Synapse quantification by live imaging (Fig 3C,D)

Animals screened for bright sparse dsRed expression were reared in D-serine or control rearing medium for 48 h (corresponding to Day 2 of daily imaging). Animals were imaged with the same setup as for daily imaging. Z-series optical sections were collected at 1 µm intervals using Fluoview software (version 5.0) for both the red (dsRed) and green (PSD95-GFP) channels. All image z-stacks were processed with FIJI software using the background subtraction (rolling ball 10 µm) and median (radius 2 µm) filtering functions. Images were made binary using MaxEntropy and Moments auto-thresholding for the dsRed and PSD95-GFP channels, respectively. To identify synapses, the Analyze Particles function with a size criterion of 0.1–5.0 μm^2^ was applied to identify synaptic puncta marked with PSD95-GFP. The logical AND of the two channels, with the size criterion of 0.1–5.0 μm^2^ area was again applied to produce an estimate of synaptic puncta density over the dendritic volume. Analysis was performed blind to experimental condition.

### In vivo two-photon calcium imaging

Animals were imaged with a Olympus XLUMPlanFLN 20X water-immersion objective (1.0 NA) mounted on a commercial high-speed resonance scanner-based multiphoton microscope (Thorlabs) with piezoelectric objective focusing (Physik Instrumente). To detect GCaMP6s, an excitation wavelength of 910 nm was used, and emission signal was collected through a 525/50 nm bandpass filter. A 250 μm square image field captured the full tectal neuropil, and images were collected from a single optical section at 15 Hz, or a stack of 4 optical sections at ∼6 Hz, sufficient to achieve the Nyquist criterion for signal detection given the long half-life of GCaMPs activation (Chen et al., 2013). Calcium recordings were corrected for *xy* drift using NoRMCorre (Pnevmatikakis and Giovannucci, 2017), and analyzed using custom Matlab and Python scripts.

#### Calcium response decay in presence of MK-801 (Fig 3-1)

Transgenic tadpoles screened for bright GCaMP6s expression were reared in D-serine or control media for 48 hr at stage 48 (corresponding to the stage at Day 2 of daily imaging). Animals were immobilized with 2 mM pancuronium bromide (Tocris). To ensure rapid drug delivery to the optic tectum, a small incision was made with a 30 ga syringe needle on the dorsal surface of the animal’s skin at the midline, exposing the ventricle underneath (Benfey et al., 2020, bioRxiv). The tadpole was then embedded in 0.8% w/v UltraPureTM low melting point agarose (Thermo Fisher) on a 6 cm petri dish lid, gently brushing away excess agarose from the skin incision on the dorsal surface. The petri dish was filled with 9 mL of 0.1x MBSH, and animals were allowed to stabilize in the imaging set-up for 20 min in darkness.

Immediately prior to imaging, 10 μM MK-801 was delivered by dissolving it in 1mL of 0.1x MBSH and using a micropipette to introduce the drug into the bath. During imaging, the animal was presented with 10 ms full-field light flashes at 0.0625 Hz from a red luxeon LED controlled by a UnoR3 board and a custom Matlab script. This 0.0625 Hz stroboscopic stimulation was selected to prevent temporal frequency adaptation during the imaging period, since we previously characterized that it elicits robust calcium activity in RGC axon terminals with no discernible rundown in the absence of drug application (Chorghay et al., 2021, bioRxiv). Each imaging session lasted 15 mins to image GCaMP6s activity to 0.0625 Hz in the presence of the use-dependent NMDAR antagonist MK-801.

To estimate the tectal neuropil response to each subsequent stimulus presentation (LED flash), the raw continuous calcium fluorescence trace was first smoothed (Gaussian filter, sigma = 4 frames) to identify local minima. The raw trace was then split into several segments, each segment spanning two successive local minima, corresponding to the response window for a single stimulus presentation. The neural response to individual stimulus presentation was quantified as the area under the curve for each segment of the calcium trace, divided by the width of the segment. The neural response to each stimulus presentation over the entire 15 min imaging session was thereafter fitted with an exponentially decaying function to characterize the decay time constant of the system. For each animal, two optical sections at comparable depths across animals were analyzed, consistently capturing a clear cross-section of the neuropil with sufficient GCaMP6s expression.

#### Retinotopic mapping (Fig 4)

Animals selected for bright GCaMP6s and mCherry expression restricted to one lateral half of their body were transferred to D-serine or control rearing medium from stage 37 onwards and imaged a week later at stage 48, since RGC axons first arrive at the anterior end of the tectum at stage 37 (Holt and Harris, 1983). Prior to imaging, tadpoles were immobilized with 2 mM pancuronium bromide (Tocris), and embedded in 1% low melting point agarose in a custom chamber with a glass coverslip window on one side through which the animal could view visual stimuli presented on an LED screen. A #29 Wratten filter over the LED screen prevented light from the display from interfering with detection of the calcium signal. Custom Matlab scripts based on the Psychophysics Toolbox (Brainard, 1997; Pelli, 1997; Kleiner et al., 2007) were used to generate the visual stimuli and synchronize stimulus presentation with image capture. The display spanned roughly 108° visual angle in azimuth and 84° in elevation. Each imaging session lasted for roughly 1.5 hours to characterize GCaMP6s activity in one tectal hemisphere.

The functional retinotopic map was extracted by presenting either static or drifting dark bar stimuli on the LED display, and correlating the stimulus-evoked response in the tectum to the stimulus positions. For “phase mapping”, the animal was repeatedly presented with a 18°-wide dark bar on a bright background drifting at a constant rate of approximately 10.8°/sec along the full span of the anterior-posterior or superior-inferior axis of the screen. A Fourier transform was performed on response traces, with the Fourier phase and amplitude component at the frequency of stimulus presentation revealing the peak stimulus-evoked response. This was calculated for every pixel to create a phase map of peak stimulus-evoked responses, which converts to a representation of visual space in the tectum (Kalatsky and Stryker, 2003). The signal-to-noise ratios of responses were calculated as Ar/σ, where Ar is amplitude at the stimulus frequency, and σ is the standard deviation of all amplitudes at frequencies above the stimulus frequency. The receptive field position is represented by the absolute phase. For each sample pixel, we calculated discontinuity as the mean distance between the absolute phase of that pixel and the phase of all its neighboring pixels within a 3.5 µm radius. Sample pixels with less than 20 valid neighbors were discarded, and tecta with less than 4000 retained sample pixels were excluded from statistical analyses. For a given optical section, we sampled 5000 pixels in the neuropil area with SNR > 1 to calculate mean “map discontinuity”.

For “grid mapping”, the animal was presented with vertical or horizontal 18°-wide dark bars at 5 equidistantly spaced positions along the azimuth or elevation axis respectively, in a randomized fashion. Then for each pixel, an optimal stimulus position was calculated based on the pixel’s weighted average ΔF/F₀ response to each stimulus position to estimate the pixel’s receptive field center. Cell body ROIs were automatically segmented using Cellpose (Stringer et al., 2021) and ΔF/F₀ responses were averaged within each ROI. We quantified “receptive field sharpness” as the average ΔF/F₀ response to the 2 or 3 stimulus positions closest to the optimal stimulus position (grid) divided by the average response to the remaining stimulus positions in the periphery. Only cell bodies with maximal stimulus response ΔF/F₀ > 2 and optimal stimulus positions falling between the 3 central stimulus positions were evaluated. Cell body segments smaller than 30 pixels and animals with less than 30 cell bodies fitting the evaluation criteria were excluded.

### Statistical analysis

Statistical analyses were performed in GraphPad Prism 8.0. Normality of the data distributions was confirmed using the Shapiro−Wilk test. The details for the statistical tests for each experiment can be found in the figure legend and figures. All quantification is graphed as mean ± SEM, unless indicated.

## Results

### Retinotectal postsynaptic arbors are stabilized by D-serine exposure

To visualize dendritic arbor morphology of neurons in the optic tectum, we performed Cre-mediated single cell labelling by electroporation (CREMSCLE) to sparsely label recently differentiated tectal cells with EGFP (Schohl et al., 2020), then performed daily interval *in vivo* two-photon imaging. After collecting a baseline image of individually labeled tectal neurons (Day 0), we reared animals in 100 μM D-serine, previously shown to significantly enhance retinotectal NMDAR currents (Van Horn et al., 2017),over the following 3 d and imaged cells every 24 h. We observed that tectal neuron dendritic arbors reared in D-serine became more compact compared to controls (Fig 1). Specifically, in the D-serine condition, there was a reduction in arbor length apparent by Day 2 (Fig 1B), as well as a lower total number of dendric branches (Fig 1C). Branch density in the dendritic arbors of animals exposed to D-serine increased over time (Fig 1D). Sholl analysis revealed that D-serine rearing shifted the distribution of branches closer to the cell body (Fig 1E). In contrast, we failed to observe significant effects of D-serine rearing on dendritic arbor growth in cells that had been labelled by single-cell electroporation of EGFP (Haas et al., 2001), (Fig 1-1), likely because this method labels both immature and mature neurons. Therefore, D-serine rearing was associated with compaction of dendritic arbors specifically in recently differentiated, immature neurons.

To investigate whether dendritic arbors are driven to stabilize prematurely by D-serine administration, we studied branch dynamics with short interval imaging every 10 min for 2 h. As daily imaging had revealed significant differences in arbor length emerging by day 2, and in order to compare with previously published axonal data (Van Horn et al., 2017), we imaged dynamic remodeling of dendritic arbors from animals that had been reared for 48 h in D-serine. While the general trend toward more compact, less branched arbors following D-serine treatment seen in the daily imaging experiments (Fig 1) was also discernible in these cells, differences in total arbor length, branch number, and rates of branch addition and loss did not reach statistical significance (Fig 2A-E). On the other hand, branch elongation and retraction lengths were significantly reduced in these D-serine-reared animals compared to controls (Fig 2F, G). Thus, D-serine appeared to stabilize dendritic morphology in these recently differentiated neurons, most notably by limiting dendritic branch elongation and retraction.

Since NMDAR activity is thought to underlie coincidence detection during patterned neuronal activity, and in turn, to contribute to downstream structural plasticity, we also asked whether the postsynaptic arbor dynamics were affected by visual stimulation. During the 2 h of short interval imaging, animals reared in D-serine or control medium experienced 1 h of darkness followed by 1 hr of visual stimulation consisting of 1 Hz stroboscopic flashes (“strobe”), which promotes synchronous coactivation of RGC axons. There was a significant shift in the dynamics of the dendritic arbor away from net addition of branches (ratio of added-to-lost > 1) during darkness to more equal rates of addition and loss under strobe (Fig 2H), as well as a decrease in mean arbor length change per 10 min, averaged for strobe versus darkness (Fig 2I), both of which are consistent with increased branch stability during synchronous stimulation. Hence, overall postsynaptic arbor growth and elaboration were stabilized under strobe, but due to the absence of a statistical interaction in our dataset, it does not necessarily support the idea that D-serine might enhance the experience-dependent stabilization effect.

### Retinotectal synaptic density is increased with D-serine exposure

Previous reports from our lab indicated that NMDAR signal enhancement by D-serine rearing primarily drives the maturation of NMDAR-only “silent” synapses, as revealed by an increase in frequency as opposed to amplitude of unitary synaptic responses (Van Horn et al., 2017). These electrophysiological measurements could not, however, fully rule out the formation of new synapses. In light of our finding that dendritic morphogenesis was affected by D-serine rearing, we examined the effects of chronic D-serine on synapse density using anatomical measures. Immunohistochemical labelling of presynaptic (SV2) and postsynaptic (GluA1) markers on cryostat sections of optic tectum in animals reared in D-serine for 48 h was performed, enabling quantification of “anatomical synapses’’ based on their apparent colocalization in confocal micrographs, as described in Sild et al. (2016). Anatomical synapse density was significantly increased in the D-serine tectal hemispheres (Fig 3A-B).

Given our observation that the dendritic arbor stabilized in recently differentiated tectal cells, we further asked if synaptic density changes were visible in the dendritic arbors of these cells. To measure this, we electroporated the optic tectum to express Post-Synaptic Density protein 95 fused to EGFP (PSD95-GFP) along with sparse single-cell expression of dsRed by CREMSCLE. The PSD95-GFP punctum density in these dsRed-labelled neurons was then quantified. We found significantly higher PSD95 punctum density in the dendritic arbors of these recently differentiated tectal neurons in animals reared in D-serine compared to control animals (Fig 3C-D). Taken together, these data suggest that D-serine treatment results in higher synapse density.

It has been reported that chronic activation of the co-agonist site of the NMDAR by D-serine can prime receptor internalization in hippocampal neurons (Nong et al., 2003). To assess whether D-serine exposure resulted in a loss of NMDAR functionality, we compared the NMDAR-dependent component of visually-evoked responses in GCaMP6s transgenic tadpoles reared for 48 h in D-serine or control medium (Fig 3-1). The NMDAR contribution to the visually-evoked calcium response in the optic tectum was estimated by measuring the integrated tectal neuropil response to visual stimulation in the presence of the use-dependent NMDAR channel blocker MK-801. Because MK-801 irreversibly blocks open channels, it is expected to incrementally decrease the NMDAR-mediated component of the calcium response with each sequential stimulus presentation. We calculated the exponential rates of decay of the visually-evoked calcium responses, and found no significant difference in the MK-801-mediated decay time constants between the two conditions. If anything, the rate of decay was slightly but non-significantly greater in the D-serine reared animals, consistent with the longer NMDAR channel opening expected in the presence of high co-agonist levels (Martina et al., 2003). This suggests that NMDARs continued to contribute robustly to retinotectal synaptic input, even following prolonged chronic D-serine exposure.

**Figure 3-1.**
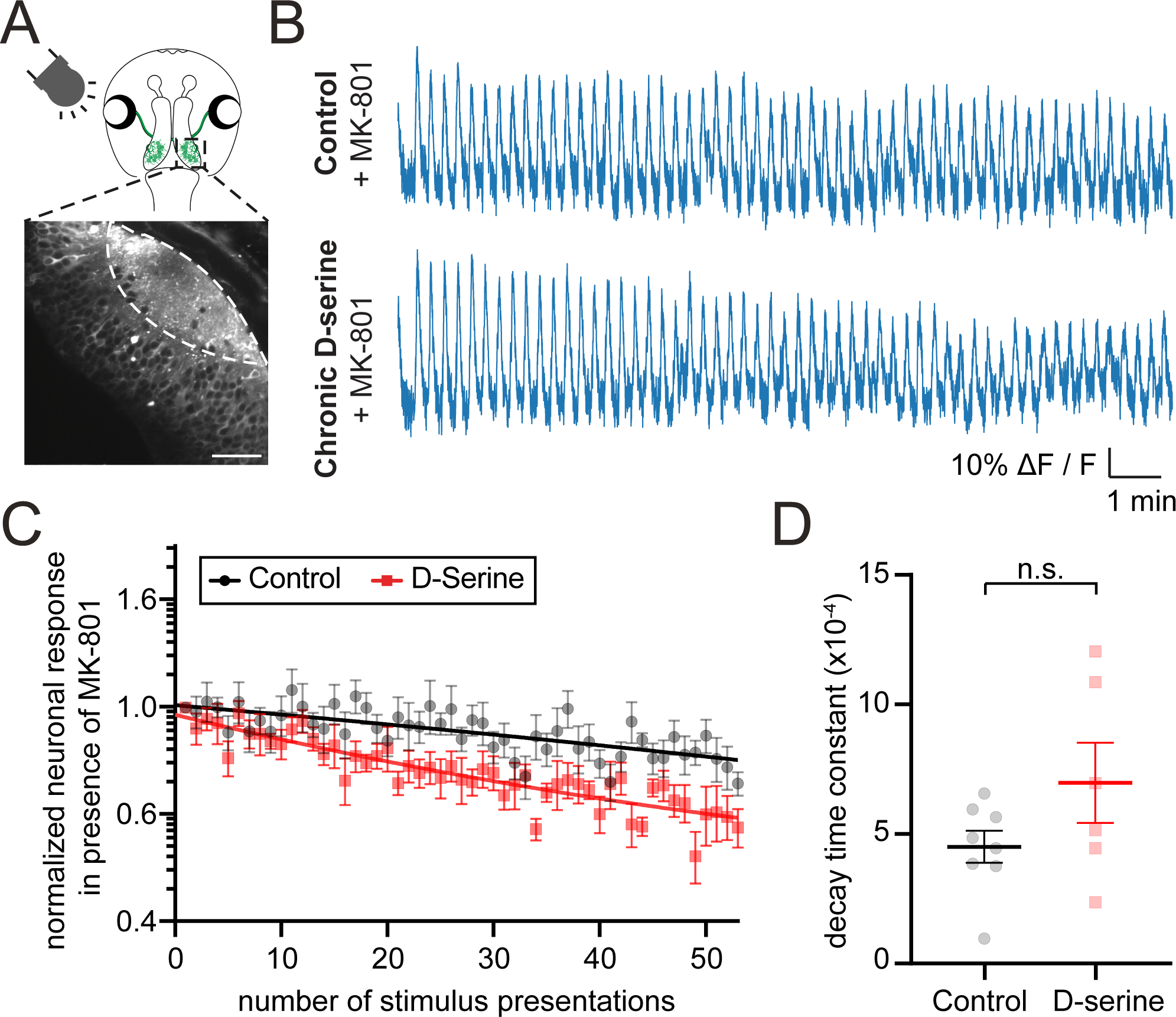
MK-801-induced decay of tectal neuropil responses in animals reared in D-serine for 48 h. (A) Calcium fluorescence in the tectal neuropil of GCaMP6s transgenic tadpoles undergoing 0.0625 Hz stroboscopic stimulation was observed by two-photon microscopy. Scale bar: 50 µm. (B) Representative tectal neuropil calcium responses to 0.0625 Hz strobe in the presence of MK-801 over a 15 min imaging period from animals reared in D-serine or control medium. (C) D-serine and control group-averaged neuronal responses to each stimulus presentation, normalised to the initial stimulus-evoked neuronal response, show a decay in the presence of MK-801. Dark lines: exponential fit of the response curve (semi-log plot). (D) Decay time constant shows no significant differences across conditions. [t-test, 2 optical sections per animal from n=3 animals for D-serine, n=4 for control.]

### Retinotectal map refinement is altered by D-serine rearing

Our group previously demonstrated using whole cell synaptic current recordings *in vivo* that rearing tadpoles in D-serine enlarged the visual input receptive fields in response to incremental (light-ON) visual flashes (Van Horn et al., 2017). To investigate how D-serine rearing may alter properties of the retinotopic map, we used calcium imaging to extract functional topographic maps from the optic tecta of tadpoles reared in D-serine or control media. We performed mRNA injections into one blastomere at the two-cell stage of development to generate bilateral mosaic tadpoles expressing GCaMP6s on one side of the animal, thus labelling tectal cells but not the RGC axons innervating them from the contralateral eye. We allowed these animals to develop and then performed two-photon calcium imaging of the optic tectum while presenting receptive field mapping stimuli (Fig 4A). Animals were imaged at stage 48 after being reared in D-serine from stage 37, corresponding to the developmental stage at which RGC axons first arrive to innervate the tectum (Holt and Harris, 1983).

To extract retinotopic maps in the optic tectum, tadpoles were repeatedly presented with a slowly moving dark bar on a bright background drifting continuously along the azimuth (anterior-posterior) or elevation (superior-inferior) axes. A Fourier analysis of GCaMP fluorescence responses revealed the stimulus phase, corresponding to the bar position, at which the optimal response occurred (Kalatsky and Stryker, 2003). This was calculated for each pixel to reveal the retinotopic map in the whole tectum (Fig 4B). We used these phase maps of the neuropil responses to quantify “map discontinuity” as the mean distance between receptive field positions of neighboring pixels, such that low mean discontinuity indicates a smooth map. We detected a slight but non-significant reduction in the map discontinuity of D-serine animals along the azimuth axis (Fig 4C) and no difference in the elevation axis (Fig 4D). Importantly, D-serine rearing did not disrupt the overall coarse topographic organization of the retinotectal maps.

To measure the response properties of individual tectal neurons, tadpoles were presented with a dark bar on a bright background repeatedly appearing in random order at one of 5 positions along the azimuth or elevation axes. Individual cell somata were segmented from the two-photon images and the receptive field position for each tectal cell body was estimated based on the relative strength of the responses to the stimuli in each of the 5 positions (grid mapping; Fig 4E, F). “Receptive field sharpness” was defined as the average fluorescence change to stimuli presented in the optimal stimulus positions, divided by the mean response to stimulation of the remaining non-optimal stimulus positions. Receptive field sharpness in the azimuth (Fig 4G) and elevation (Fig 4H) axes increased following D-serine rearing, as indicated by a shift in the cumulative probability distributions of sharpness measurements for all cells toward higher values. Thus, our data suggest that D-serine rearing leads to sharper and therefore more compact visual receptive fields for tectal neurons, consistent with arbor compaction and increased synapse density of tectal cell dendrites.

## Discussion

Our study uses chronic administration of the NMDAR co-agonist D-serine to reveal morphological changes to the retinotectal circuit, including increased dendritic arbor compaction and reduced branch dynamics in recently differentiated neurons, as well as overall higher synapse density in the tectal neuropil. Functionally, calcium imaging of the optic tectum showed increased receptive field sharpness of tectal neurons raised in D-serine. Together, these findings suggest that the availability of D-serine at glutamatergic synapses influences refinement of the developing retinotectal circuit, confirming the importance of NMDARs in developmental plasticity.

Our morphological observations of decreased dendritic growth under conditions of NMDAR enhancement by D-serine administration were initially surprising given earlier reports that NMDAR blockade with APV *in vivo* also reduced dendritic growth over 24 h (Rajan and Cline, 1998) and decreased dendritic branch size and dynamics over 2 h of imaging (Rajan et al., 1999). However, blockade using APV may not provide the complete picture with regards to NMDAR function as NMDARs may perform different roles in different cell types and cellular compartments. For example, GluN1 knockdown in postsynaptic tectal cells leads to fewer dendritic branches compared to cells in which GluN1 is knocked down in their presynaptic RGC inputs (Kesner et al., 2020). To interpret these findings on whether increased NMDAR activation leads to enhanced stability of the postsynaptic arbor, it is necessary to consider the distinct effects on the circuit of different NMDAR manipulations, including systemic or targeted, acute or chronic, receptor knockdown or drug administration.

Here, we report that enhancing NMDAR activity with D-serine for 48 h led to compaction of the dendritic arbor and attenuated dynamic branch elaboration and retraction specifically in recently differentiated neurons. Our results are in line with experiments showing that the overexpression of CaMKIIa, a key effector downstream of the NMDAR, also slows dendritic growth of immature but not of more mature neurons, whereas CaMKII inhibition increases the dendritic growth only of mature neurons (Wu and Cline, 1998). Thus, NMDAR activity appears to limit growth and stabilize arbors of more immature neurons, perhaps acting as an activity-dependent mediator of neuronal maturation.

To further understand how coincidence detection by NMDARs converts neuronal activity into cues for topographic refinement, one approach has been to manipulate sensory visual experience in order to induce patterned neuronal activity and study its effects in real time (Kutsarova et al., 2016). Sequentially activating LED arrays to mimic visual flow promotes dendritic arbor growth (Sin et al., 2002). Stroboscopic visual stimulation of both eyes for synchronous coactivation of RGC inputs stabilizes axonal arbors (Munz et al., 2014), while asynchronous activation is accompanied by increased arbor dynamics (Rahman et al., 2020). NMDAR blockade mitigates the stabilizing effects of visual stimulation on arbor morphology (Sin et al., 2002; Munz et al., 2014). Here, we observed that strobe stimuli shifted the dynamics of dendritic arbor growth to favor matched rates of growth and retraction. In this way, synchronous coactivation of the visual field may restrict net dendritic arbor elaboration. However, surprisingly, D-serine rearing did not appear to significantly enhance this activity-dependent change in dynamics. This finding may indicate that the influence of chronic D-serine exposure on the circuit acts over much longer timescales than some other aspects of dendritic branch dynamics.

The shift from small, simple arbors to larger, more complex arbors as the neuron matures also corresponds to the well-characterized shift from predominantly NMDAR-mediated glutamatergic neurotransmission typical of immature synapses to both AMPAR- and NMDAR-mediated neurotransmission at mature synapses (Wu et al., 1996; Petralia et al., 1999). Cells with shorter dendritic arbors (<200 μm length) tend to show an AMPA/NMDA ratio <1, whereas those with larger arbors (>200 μm length) have an AMPA/NMDA ratio >1 (Rajan and Cline, 1998). Indeed, as proposed in Vaughn’s synaptotropic hypothesis, branch stability may be conferred by the presence of a stable synapse: synapses marked with GFP-VAMP2 or synaptophysin-GFP act as the sites of new axonal branch tip formation (Alsina et al., 2001; Meyer and Smith, 2006; Ruthazer et al., 2006); stable synapses are correlated with stabilised axonal branches (Vaughn et al., 1989; Meyer and Smith, 2006; Ruthazer et al., 2006; Cline and Haas, 2008); and synapse maturation versus disassembly has been linked to dendritic arbor elaboration versus retraction (Niell et al., 2004; Haas et al., 2006; Chen et al., 2010). Previously, our group reported that enhanced NMDAR activation with D-serine administration increased spontaneous miniature EPSC frequencies and decreased paired pulse ratios consistent with an increased probability of release (Van Horn et al., 2017). Increased AMPA/NMDA ratios were also observed in D-serine reared animals, further inferring the maturation of immature NMDAR-only “silent” synapses (Van Horn et al., 2017). In the anatomical measurements of synapse density of tectal neurons that we report here, increased synapse density was observed following D-serine exposure, further supporting the idea that D-serine promotes a stabilization phenotype.

These NMDAR-related changes in morphology have been tied to the development and maintenance of retinotopic maps. Chronic NMDAR blockade leads to a degradation of RGC projection convergence within the optic tectum (Cline and Constantine-Paton, 1989) and to desegregation of ocular dominance bands in dually innervated tecta in amphibians (Cline and Constantine-Paton, 1990; Ruthazer et al., 2003). NMDAR blockade also leads to a functional enlargement of visual receptive fields in the tectum (Dong et al., 2009). Finally, receptive field sizes for ON responses increase after 48 h of D-serine exposure (Van Horn et al., 2017), while the ratio of ON/OFF response strength increases with GluN1 postsynaptic knockdown (Kesner et al., 2020). Overall, these lines of evidence suggest that NMDAR signaling is involved in the targeting and growth of RGC inputs, strengthening of appropriate synapses, and ultimately, topographic map refinement. Examining tectal receptive field properties by calcium imaging, we saw that raising animals in D-serine increased receptive field sharpness of individual neurons but did not significantly affect the continuity of the full topographic map in the tectal neuropil. Given that the coarse organization of the retinotopic map is largely preserved under manipulations of NMDAR activity that alter receptive field size and sharpness, we suggest that NMDAR enhancement by D-serine exposure fails to grossly disrupt the overall topographic organization established by molecular guidance cues (Higenell et al., 2012; Triplett & Feldheim, 2012) but may modulate synaptic plasticity, leading to subtle local alterations in the retinotopic map.

Consistent with our observations in the tadpole visual system, in other model systems, such as in the visual thalamus of the ferret, dendritic arbor size and spine density were increased in response to one week of APV administration (Rocha and Sur, 1995). In the mammalian homologue of the tectum, the superior colliculus, NMDAR blockade leads to less precise innervation by RGC axons (Simon et al., 1992) and enlarged receptive fields (Huang and Pallas, 2001). Furthermore, chronic inhibition of GluN1 translation using antisense technology during development prevents the development of orientation selectivity in the ferret primary visual cortex (Ramoa et al., 2001). In the somatosensory system of mice with GluN1 knockdown, barrelette cells receiving trigeminal whisker afferents develop longer dendrites with no orientation bias (Lee et al., 2005a). In turn, these animals, as well as mice with cortex-specific GluN1 knockdown, show disrupted branching patterns of the whisker afferents (Lee et al., 2005a; Lee et al., 2005b). Loss of functional NMDARs is crucial for somatosensory patterning, both at the level of the brainstem (Iwasato et al., 1997) and the cortex (Zhou et al., 2020).

Finally, our study involved the application of exogenous D-serine to better understand NMDAR-mediated synaptic plasticity. However, it will be important to investigate the contributions of endogenously released D-serine under both physiological and pathological conditions. Since D-serine synthesis and release is thought to be regulated in concert by neurons (Wolosker et al., 2017) and glia, particularly astrocytic glia (Papouin et al., 2017), this suggests a metaplasticity mechanism wherein astrocyte-regulated D-serine availability at glutamatergic synapses shapes NMDAR-mediated plasticity. Aberrant changes in NMDAR activation and D-serine synthesis and availability have been implicated in neuroinflammation and excitotoxicity in Alzheimer’s disease, amyotrophic sclerosis, ischemia, and schizophrenia (Pollegioni and Sacchi, 2010; Van Horn et al., 2013). It has been suggested that pathology shifts the primary source of D-serine from neurons to glia, and clinical trials targeting D-serine as well as its biosynthetic and regulatory pathways are currently underway for therapeutic use (Ivanov and Mothet, 2019; Coyle et al., 2020). Understanding the signaling pathways and precise mechanisms by which D-serine affects structure and function will provide important insights into experience-dependent NMDAR-mediated plasticity in both health and disease.

## Conflict of Interest statement

The authors declare no competing interests.

## Acknowledgments

Confocal imaging was performed at the Neuro Microscopy Core Facility. This work was funded by a chaire de recherche from the Fonds de recherche du Québec - Santé (FRQS Grant 31036) and a Foundation Grant from the Canadian Institutes of Health Research (CIHR FDN-143238) to ER, and McGill University Integrated Program in Neuroscience Studentships to ZC and VJL.

## Notes

### Competing Interest Statement

The authors have declared no competing interest.

